# Integrative haplotype estimation with sub-linear complexity

**DOI:** 10.1101/493403

**Authors:** Olivier Delaneau, Jean-François Zagury, Matthew Robinson, Jonathan Marchini, Emmanouil Dermitzakis

**Affiliations:** Department of Computational Biology, University of Lausanne, Lausanne, Switzerland.; Department of Genetic Medicine and Development, University of Geneva, Geneva, Switzerland; Swiss Institute of Bioinformatics (SIB), University of Geneva, Geneva, Switzerland.; Institute of Genetics and Genomics in Geneva, University of Geneva, Geneva, Switzerland.; Chaire de Bioinformatique, Conservatoire National des Arts et Métiers, Paris, France.; Department of Statistics, University of Oxford, UK.

## Abstract

The number of human genomes being genotyped or sequenced increases exponentially and efficient haplotype estimation methods able to handle this amount of data are now required. Here, we present a new method, SHAPEIT4, which substantially improves upon other methods to process large genotype and high coverage sequencing datasets. It notably exhibits sub-linear scaling with sample size, provides highly accurate haplotypes and allows integrating external phasing information such as large reference panels of haplotypes, collections of pre-phased variants and long sequencing reads. We provide SHAPET4 in an open source format on https://odelaneau.github.io/shapeit4/ and demonstrate its performance in terms of accuracy and running times on two gold standard datasets: the UK Biobank data and the Genome In A Bottle.

## Introduction

Haplotypes are key features of disease and population genetic analyses^1^ and data sets in which they prove to be useful evolve in two main directions. On the one hand, cost reduction of SNP arrays allow genotyping hundreds of thousands of individuals resulting in data sets such as the UK Biobank (UKB) that regroup genotype data for half a million samples^2^. To efficiently estimate haplotypes on this scale, three methods, eagle2^3,4^, shapeit3^5^ and beagle5^6,7^, have been recently proposed with running times that are either linear or close-to-linear with sample size. Data sets consisting of millions of samples are now being generated by projects such as the Million Veteran Program^8^ or by commercial companies such as 23andMe that has now genotyped more than 5 million customers so far^9^. In this context, it is unclear if the scaling with sample size offered by these methods is able to conveniently process genotype data on that scale. On the other hand, high throughput sequencing now enables exhaustive screening of millions of genetic variants within tens of thousands of individuals such as in the Haplotype Reference Consortium data set^10^. Latter developments in sequencing have also witnessed the introduction of long reads technologies such as PacBio^11^ or Oxford Nanopore^12^. By covering multiple nearby heterozygous variants in an individual, the long reads generated by these sequencing technologies allow resolving haplotypes across hundreds of kilobases. To achieve this task, commonly called haplotype assembly, multiple methods have been proposed so far such as WhatsHap^13^, HapCut^14^ or longRanger^15^. While these approaches are extremely efficient to resolve the phase between nearby variants, they do not allow full haplotype resolution across entire chromosome suggesting that additional computational steps at the population level are still required. At this point, it becomes clear that haplotype estimation is now facing two main challenges: computational efficiently to accurately process large scale data sets and data integration to exploit large reference panels of haplotypes and long sequencing reads. In this paper, we describe and benchmark a new method for haplotype estimation, SHAPEIT4, which proposes efficient solutions to these two challenges. Specifically, it allows processing either SNP array or sequencing data accurately with running times that are sub-linear with sample size, therefore making it well suited for very large scale data sets. In addition, it also facilitates the integration of additional phasing information such as reference haplotypes, long sequencing reads and sets of pre-phased variants altogether to boost the quality of the resulting haplotypes. To achieve this, the method builds on three main components: (i) the Li and Stephens model^16^ to capture long range haplotype sharing between individuals, (ii) the Positional Burrow-Wheeler Transform (PBWT)^17^ to speed up the computations involved in the Li and Stephens model and (iii) the compact representation of the solution space built in previous versions of SHAPEIT^18,19^ that allows efficient haplotype sampling and easy integration of additional phasing information^20,21^. To demonstrate its performance, we benchmark it on two gold standard data sets: the UK Biobank^2^ to evaluate its ability to process large scale SNP array data sets and on the Genome In A Bottle (GIAB)^22^ to assess its ability to leverage long sequencing read information.

## Results

### Overview of SHAPEIT4

SHAPEIT4 improves upon previous SHAPEIT versions at two main levels. As a first major improvement, it now uses an approach based on the Positional Burrow Wheeler Transform^17^ to quickly assemble small sets of informative haplotypes to condition on when estimating haplotypes. This provides a computationally efficient alternative to the previous approach based on Hamming distance^19,23^. A PBWT of haplotypes is a data structure in which any two haplotypes sharing a long prefix (i.e. match) at a given position are sorted next to each other at that position. SHAPEIT4 takes advantage of this by maintaining a PBWT of all the haplotype estimates so that long matches between haplotypes can be identified in constant time. In practice, SHAPEIT4 works within overlapping genomic regions (of 2Mb by default) and proceeds as follows to update the phase of an individual in a given region: (i) it interrogates the PBWT arrays every 8 variants to get the P haplotypes that share the longest prefixes with the current haplotype estimates at that position, (ii) it collapses the haplotypes identified across the entire region into a list of K distinct haplotypes and (iii) it runs the Li and Stephens model conditioning on the K haplotypes (Figure 1A-B). In this approach, P is the main parameter controlling the trade-off between speed and accuracy and gives a model in which K varies and adapts to the data and region being processed as opposed to previous methods in which K is usually fixed^19,23^. Indeed, K varies depending on the length of the matches found in the PBWT: longer matches involve smaller K which typically occurs as the algorithm converges, as the level of relatedness between individuals increases and more importantly as the number of samples in the dataset increases. The latter implies, and we later show, that SHAPEIT4 scales sub-linearly with sample size. All other methods proposed so far exhibit at best linear or close-to-linear scaling. In other words, the time spent per genome decreases as the total number of genomes being processed increases.

**Figure 1:**
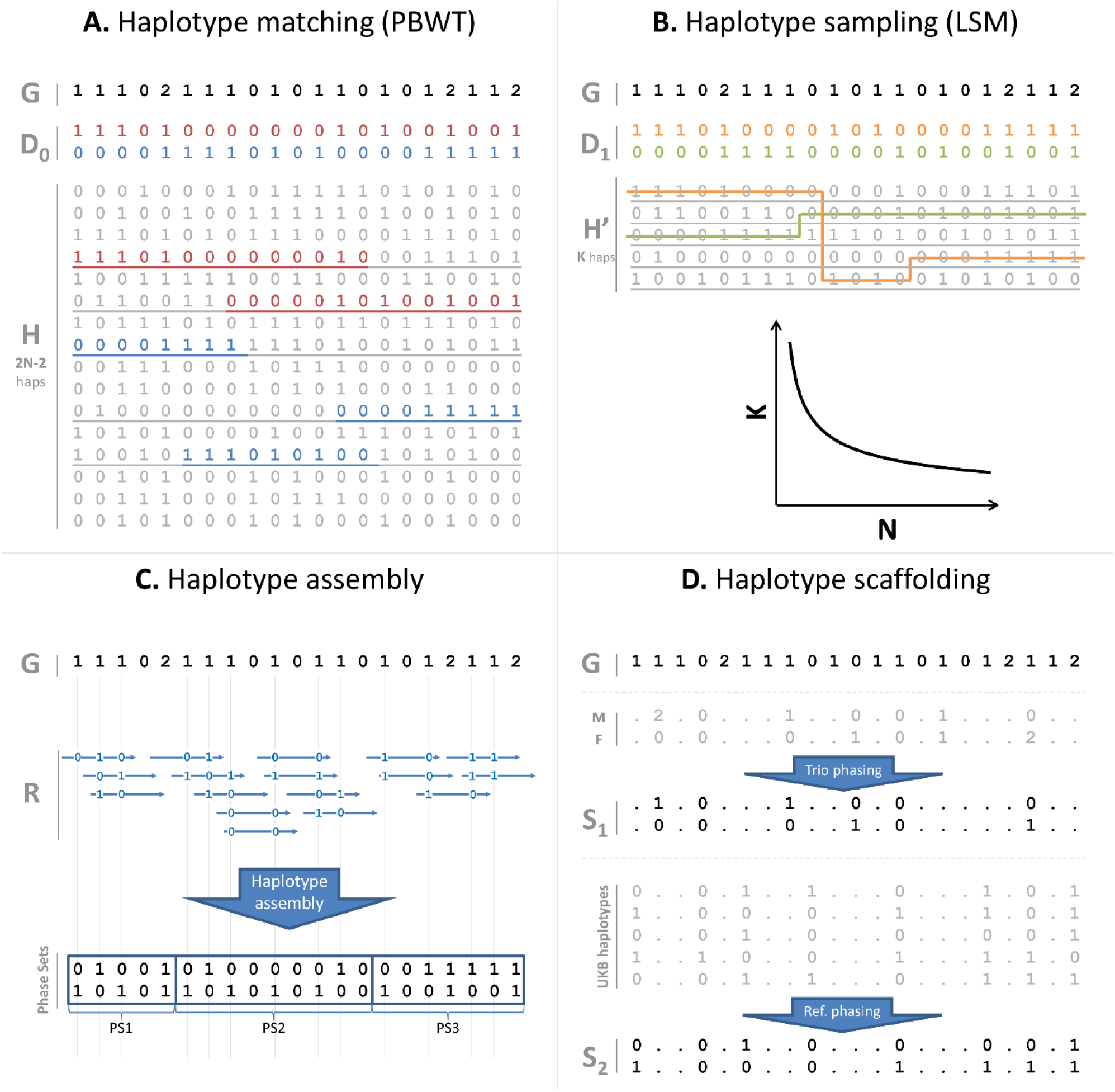
SHAPEIT4 overview. In all panels is shown the unphased genotype data for individual G. (**A**) Selection of a small number of informative haplotypes. Conditioning haplotypes with long matches with the current estimate D_0_ identified by PBWT are underlined. Matches are shown in red and blue depending on the matched haplotype. (**B**) Illustration of the Li & Stephens model run on the 5 informative haplotypes with long matches identified with PBWT. This gives new estimates D_1_ for G. (**C**) An example of a set R of phase informative reads (i.e. overlapping multiple heterozygous genotypes) for individual G. Haplotype assembly on R gives three phase sets (PS1, PS2 and PS3). The phase sets are the information used by SHAPEIT4 when estimating the haplotypes for G. (**D**) Two examples of haplotype scaffolds for G. S_1_ is derived from trio information (M and F are genotype data for the mother and father of G). S_2_ is derived from a reference panel such as UKB. Only variants in G in the overlap with UKB are phased using all UKB haplotypes as reference panel.

As a second improvement, SHAPEIT4 offers the possibility to integrate three additional layers of phasing information when estimating haplotypes: a reference panel of haplotypes, phase information contained in sequencing reads and subsets of genotypes at which the phase is known a priori (termed as haplotype scaffold). This builds on previous work developed as part of SHAPEIT2^20,21^ so that all these layers of information can be conveniently and simultaneously used. Adapting the method to leverage reference panels of haplotypes is straightforward: the PBWT matching procedure was just extended to also consider the reference haplotypes when selecting conditioning haplotypes. Concerning the two other layers of information, we implemented them as constraints in the haplotype sampling scheme. Specifically, we leverage phase information contained in sequencing reads in a two steps approach. First, we perform haplotype assembly with methods such as WhatsHap for instance^13^. This essentially regroups nearby heterozygous genotypes into phase sets when they are overlapped by the same sequencing reads (Figure 1C). Then, we model the resulting phase sets as probabilistic constraints in the SHAPEIT4 haplotype sampling scheme so that haplotype configurations consistent with them are favoured but not necessarily sampled. This is controlled by a parameter that defines the expected error rates in the phase sets (default is 0.0001). As a consequence, depending on the certainty of the population based phasing calls, we basically have two possible scenarios: (i) uncertain calls can get informed by the phase sets which typically occur at rare variants and (ii) calls with high certainty can correct phase sets when they contain errors. For the haplotype scaffold, we explored two possible strategies in this work: a family based scaffold that we derived from genotyped parents and a population based scaffold that we derived from very large reference panels of haplotypes (Figure 1D). In both cases, this gives reliable haplotype estimates defined at a sparser set of variants that we leverage by enforcing SHAPEIT4 to only sample haplotypes that are fully consistent with the available haplotype scaffolds. This helps the algorithm to converge towards good resolutions by pruning out unlikely configurations.

### Phasing large scale datasets

We assessed the performance of SHAPEIT4 on genotype data sets with large sample sizes and compared it to the following methods: SHAPEIT3^5^, Eagle2^3,4^ and Beagle5^6,7^. To do so, we built multiple subsets of the UKB data set comprising up to ˜400,000 individuals, 500 of them being trio children for whom haplotypes can be derived with certainty from the family information. For each phasing scenario, we measured the overall running times and the switch error rates on the 500 trio children. Overall, we find that all tested methods provide haplotype estimates with low error rates that substantially decrease as sample size increases (Figure 2A). A closer look reveals that both Beagle5 and SHAPEIT4-P=4 significantly outperforms all other methods on all tested configurations. For instance, on the largest sample size, we get the following ranking: SHAPEIT4-P=4 (0.117%), Beagle5 (0.125%), SHAPEIT4-P=2 (0.139%), Eagle2 (0.178%), SHAPEIT4-P=1 (0.202%) and SHAPEIT3 (0.356%). As expected, increasing P yields to appreciable improvements in accuracy: for instance, the P=2 and P=4 configurations decrease the error rate of P=1 by 31% and 42% on the largest sample size, respectively. This shows that having multiple candidates to copy from at a given position helps the algorithm to reach good estimates. Accuracy of the haplotype estimates can also be assessed by looking at the mean length of haplotype segments free of any switch errors. These segments become very long when phasing 400,000 samples: 15.50Mb and 14.75Mb for SHAPEIT4-P=4 and Beagle5, respectively (Supplementary Figure 1). In terms of running times, we find SHAPEIT4 to be substantially faster than all other methods, no matter the sample size (Figure 2B). For instance, SHAPEIT4-P=4 is 1.6 to 3.6 times faster than Beagle5, 1.9 to 5.8 times faster than Eagle2 and 4.1 to 11 times faster than SHAPEIT3 when phasing from 10,000 to 400,000 individuals. The speedup gets better with sample size as a consequence of sub-linear scaling. Indeed, SHAPEIT4 spends less time per genome on larger sample sizes conversely to all other methods (Figure 2C). The sublinear scaling can also be noted when looking at the variation of the number of conditioning states with sample sizes and iterations (Supplementary Figure 2A) or when looking at the coefficients of the fitted functions relating running times T to sample size N on this benchmark dataset (Supplementary Figure 2B). In terms of memory usage, we also find SHAPEIT4 to substantially improve upon SHAPEIT3 (3.5x decrease) and to offer performance comparable to Beagle5 and Eagle2 (Supplementary Figure 3). Overall, we find in this first benchmark that SHAPEIT4 exhibits the best trade-off between accuracy, speed and memory across all tested methods.

**Figure 2:**
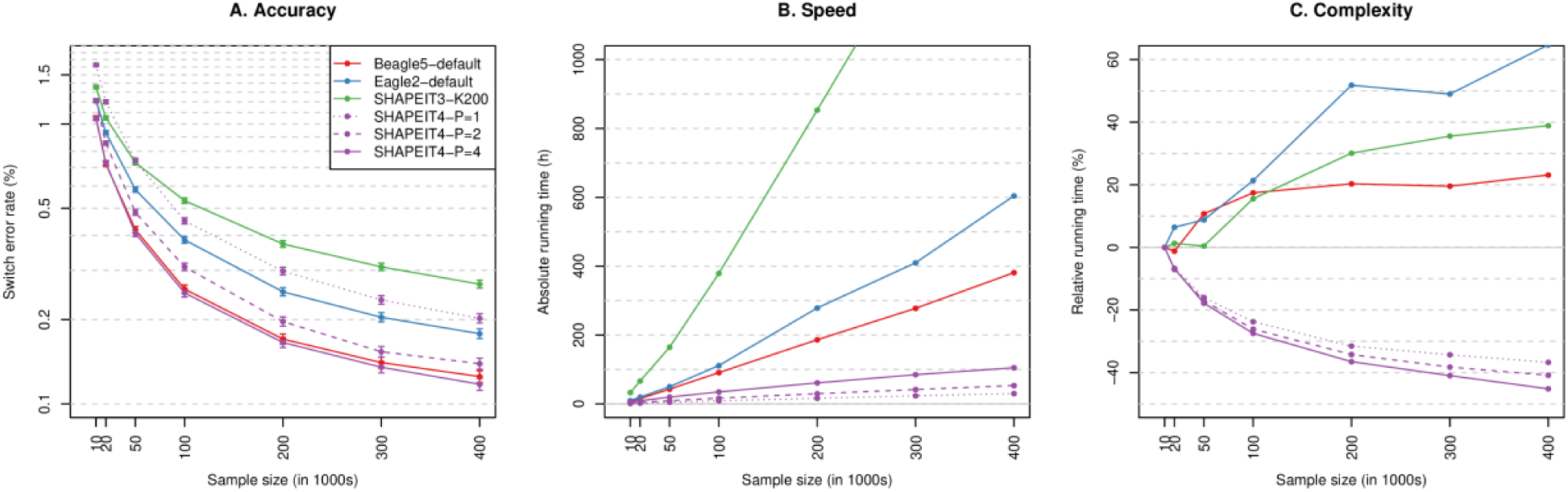
Phasing performance on large sample sizes (UKB). (**A**) Switch error rates for all tested phasing methods as a function of sample size (going from 10,000 to 400,000 individuals). For each error rate is shown the 95% binomial confidence interval. (**B**) Corresponding running times in hours measured using LINUX time command (User + System time). (**C**) Percentage variation of the running time needed to phase a single genome out of 10,000 to 400,000 genomes relative to the time needed to phase one genome out of 10,000. Positive, null or negative slopes are indicative of close-to-linear, linear or sub-linear complexities, respectively.

### Phasing from large reference panels

We also assessed the performance of SHAPEIT4 when phasing from large reference panels of haplotypes. To do so, we used the most accurate sets of haplotypes estimated for UKB as part of the first benchmark in which we removed the haplotypes of the 500 trio children. This resulted in reference panels containing from ˜1,000 to ˜800,000 reference haplotypes. We then phased against these candidate reference panels the 500 trio children alone or as part of larger datasets comprising 5,000 to 50,000 individuals, using either SHAPEIT4, Beagle5 or Eagle2. For each phasing scenario, we measured the overall running times and the switch error rates on the 500 trio children. In this second benchmark, we find the same accuracy patterns than in the first benchmark: overall error rates decrease as reference panel size increases and both SHAPEIT4-P=4 and Beagle5 outperform all other methods no matter the amount of data being processed (Figure 3A-C). In terms of running times, we find SHAPEIT4 to be faster than any other methods in all scenarios excepted one (Figure 3D-F): when phasing few samples (=500) from very large reference panels (>400,000 haplotypes) where Eagle2 seems to be slightly faster (Figure 3D). Interestingly, we also find the PBWT approach used by SHAPEIT4 to have an interesting property in this particular context: phasing large sample size sizes (=50,000 samples) is faster when using large reference panels (Figure 3F). Again here, we find SHAPEIT4 to provide overall the best trade-off between speed and accuracy when phasing from large reference panels.

**Figure 3:**
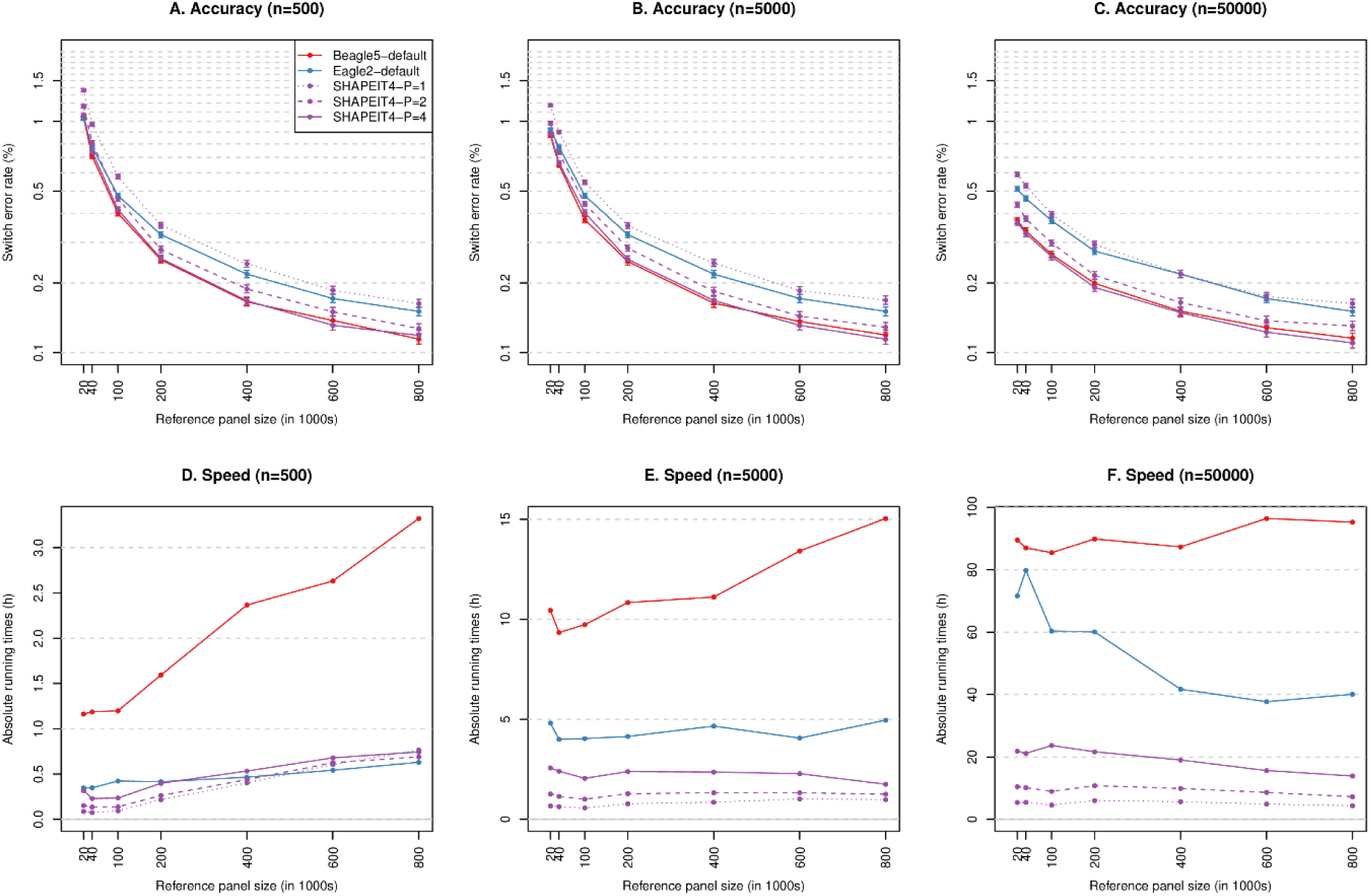
Phasing performance from large reference panels (UKB). Switch error rates (**A-C**) and running times (**D-F**) for all tested phasing methods as a function of the number of haplotypes in the reference panels. Three different sample sizes where tested for the main panel: 500 (A and D), 5000 (B and E) and 50000 (C and F). For each error rate is given the 95% binomial confidence interval.

### Phasing using sequence reads

We finally assessed the ability of SHAPEIT4 to leverage additional phase information when processing genotypes derived from high-coverage sequencing data. To do so, we merged the GIAB high coverage genotype data on chromosome 20 with 502 unrelated European individuals that have been sequenced as a part of the 1,000 Genomes project^24^. We then phased the resulting dataset using SHAPEIT4 and two different layers of phase information. First, we performed haplotype assembly (using WhatsHap; Figure 1C) for the GIAB genotype data using five types of sequencing data^22^: SOLiD5500W, Complete Genomics, Illumina HiSeq, PacBio and 10x Genomics which informed the phase at 9.8%, 35.1%, 57.9%, 90.1% and 97.7% of the heterozygous genotypes, respectively (table 1). Second, we built two different haplotype scaffolds for GIAB: (i) one derived from a larger set of samples genotyped on Illumina OMNI2.5M SNP arrays in which most heterozygous genotypes can be phased using trios and (ii) another one derived by phasing against all available UKB haplotypes (˜800,000 haplotypes; Figure 1D). Overall, (i) and (ii) allowed fixing the phase at 6.59% and 6.25% of the genotypes in the 502 individuals with sequence data, respectively. In this third benchmark, we could make the following observations. First, the error rates of SHAPEIT4 significantly decrease when using the phase information contained in the phase sets, demonstrating the ability of the methods to leverage such information (Figure 4A). Second, the various sequencing technologies exhibit large differences in terms of accuracy. Not surprisingly, technologies based on long or barcoded reads produce more accurate haplotypes with error rates as low as 0.23% and 0.07% for PacBio and 10x Genomics, respectively (Figure 4A). Third, sparse haplotype scaffolds derived either from family information or large reference panels provide appreciable boosts in accuracy, particularly at common variants (MAF>1%; Figure 4B). This demonstrates the benefit of using haplotype scaffolds when phasing sequencing data. Fourth, SHAPEIT4 allows correcting many switch errors introduced at the haplotype assembly step (Figure 4C). For instance, in our data, it corrects 26% and 79% of the switch errors obtained from Illumina HiSeq and PacBio, respectively. Finally, the value of integrating data using SHAPEIT4 can be summarized by comparing three phasing scenarios of interest (Supplementary Figure 4): (i) the default situation in which the data was directly phased without using any additional information (SER=0.82%), (ii) the situation that can often be implemented in which both Illumina HiSeq data and a UKB based haplotype scaffold are used in the estimation (SER=0.42%, 49% decrease in error rate) and (iii) the most accurate situation in which both 10x Genomics data and a trio based haplotype scaffold are used (SER=0.07%, 91% decrease in error rate).

**Table 1:** Summary statistics for GIAB sequencing data. For each type of sequencing data available for GIAB is given the total number of reads in millions, the mean length of the reads in base pairs, the mean insert size in base pairs, the means coverage and the percentage of heterozygous genotypes belonging to phase sets.

|  | #reads<br>(millions) | Mean read<br>length (bp) | Mean insert<br>size (bp) | Mean<br>coverage | %hets phased |
| --- | --- | --- | --- | --- | --- |
| **Solid5500W** | 22.13 | 68 | NA | 22 | 9.8 |
| **Complete**<br>**Genomics** | 132.6 | 33 | 282 | 77 | 35.1 |
| **Illumina HiSeq** | 13.29 | 148 | 546 | 33 | 57.9 |
| **PacBio** | 1.04 | 3707 | 2048 | 42 | 90.1 |
| **10x Genomics** | 22.27 | 88 | 194 | 33 | 97.7 |

**Figure 4:**
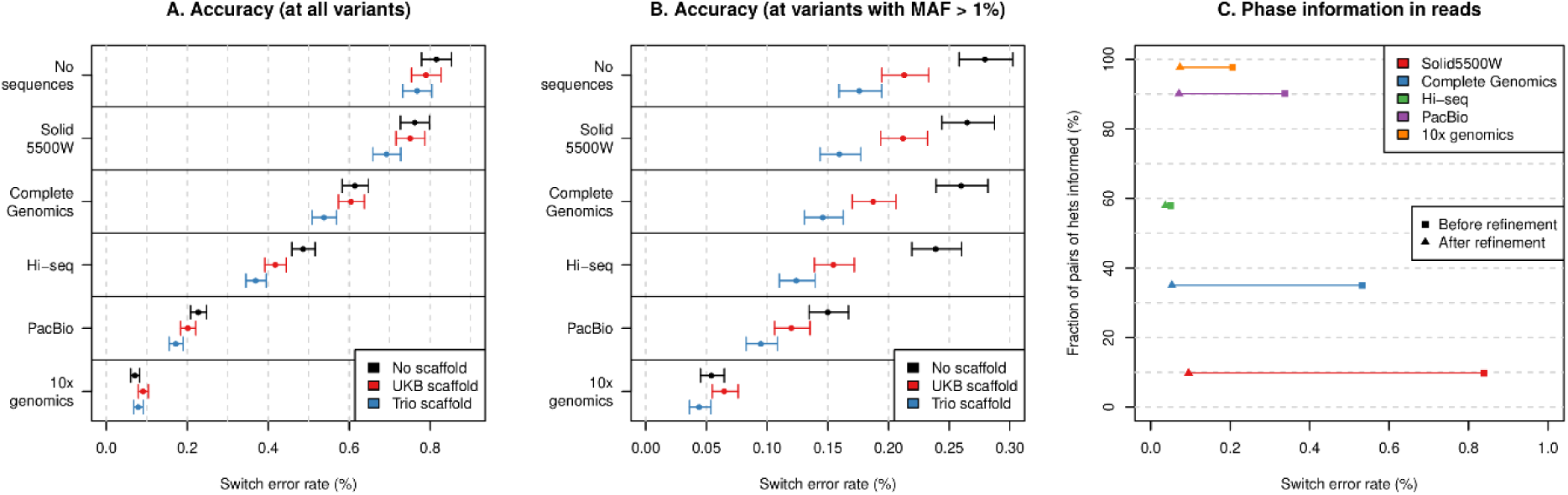
Phasing performance on high coverage sequence data (GIAB). (**A**) Switch error rates with 95% binomial confidence intervals for each combination tested of sequencing reads and haplotype scaffold. (**B**) Same information than in (A) measured only at variants with minor allele frequency (MAF) above 1%. (**C**) Switch error rates measured only at variants belonging to phase sets (i.e. haplotype assembly) before (squares) and after (triangles) refinement by SHAPEIT4.

## Discussion

We present here a new method for haplotype estimation, SHAPEIT4, that substantially improves upon existing methods in terms of speed and accuracy. One of the key improvements resides in its ability to quickly select a variable number of informative haplotypes to condition on through a PBWT based approach. The resulting method exhibits sub-linear scaling with sample size and provides highly accurate haplotype estimates for large scale data sets. We demonstrated this on the largest genotype dataset available so far, the UK Biobank, in which SHAPEIT4 offers the best trade-off between speed and accuracy across all tested methods. We predict SHAPEIT4 to be particularly well suited to process datasets reaching a million individuals and above. Of note, Eagle2^3,4^ also uses PBWT in order to build compact representations of a fixed number of conditioning haplotypes (called HapHedge structures). This fundamentally differs from our approach in which we use PBWT to identify a small and variable number of informative haplotypes to condition on. Beyond phasing, we believe this PBWT based approach to have the potential to speed up computations involved by other haplotype based models used in population genetics for admixture mapping^25^, identity-by-decent (IBD) mapping^26^ or genotype imputation^23^. Nowadays, haplotype phasing is commonly performed on imputation servers such as the UMich server^27^ based on large reference panels of haplotypes. We demonstrated that the initial implementation of SHAPEIT4 already provides high performance in this context and we predict that this can be further improved by using other PBWT algorithms (e.g. algorithm 5 from Durbin, 2014^17^). Besides computational efficiency, the SHAPEIT4 is also flexible enough to conveniently leverage additional phase information contained in long sequencing reads. We notably showed that SHAPEIT4 is able to deliver very accurate haplotypes for high coverage sequenced data when leveraging long or barcoded reads through a first step of haplotype assembly (e.g. 10x genomics). In addition, its ability to use haplotype scaffolds offers an interesting framework in which the high accuracy delivered by very large SNP array datasets can be propagated into high coverage sequencing datasets of smaller sample sizes. We predict that these two functionalities will help improving phasing servers dedicated to whole genome sequenced samples^28^. It is also important to mention here that SHAPEIT4 is not designed to perform genotype calling from low coverage sequencing and adapting the model for such data would need some additional work. Overall, we believe SHAPEIT4 to be particularly well suited to fully exploit the potential of the large datasets being generated using either SNP arrays or new sequencing technologies.

## Author contributions

OD designed the study, implemented the code, performed the experiments and wrote the manuscript. OD conceived the original ideas thanks to fruitful discussions with JM and ETD. MRR helped on the UKB data processing and the computations. JM, JFZ, MRR and ETD discussed the results with OD and contributed to the final manuscript.

## Acknowledgement

This work was conducted under UK Biobank project 35520. OD is funded by a Swiss National Science Foundation (SNSF) project grant (PP00P3_176977). MRR is funded by a Swiss National Science Foundation (SNSF) project grant (31003A-179380) and by core funding from the University of Lausanne. JM acknowledges funding for this work from the European Research Council (ERC; grant 617306). ETD is funded by the Louis-Jeantet Foundation, European Research Council and Swiss National Science Foundation.

## Online Methods

### Haplotype sampling

Let assume we have genotype data for N individuals across L variant sites that we want to phase in 2N haplotypes. To achieve this, SHAPEIT4 uses an iterative approach. It takes each individual’s genotype data G in turn and estimates a consistent pair of haplotypes D = (h_1_, h_2_) given the set H of haplotypes that has been previously estimated for the other N-1 individuals. When repeated many times, this builds a Monte Carlo Markov Chain (MCMC) is in which a single iteration consists in updating the haplotypes of all N individuals. To update the haplotypes of G, SHAPEIT4 proceeds probabilistically by sampling a new pair of haplotypes from the posterior distribution P(D|H). This posterior is based on a particular Hidden Markov Model, the Li and Stephens model (LSM^16^), that build haplotypes for G as mosaics of other haplotypes in H. This builds on the idea that individuals in a population share relatively long stretches of haplotypes inherited from common ancestors. While being accurate, the LSM is also very computationally demanding and rapidly becomes intractable on large data sets. To ameliorate this, we introduced in SHAPEIT1^18^ an algorithm able to sample from P(D|H) in linear times with the size of H. We achieved this by carrying out all LSM computations on compact graph representations of all possible haplotype configurations that are consistent with each individual (called genotype graph; see Supplementary Figure 5 for a graphical description and *Delaneau et al, 2013*^19^ for a formal one). In SHAPEIT4, we essentially use the same sampling algorithm that we completely re-implemented for better performance:

- We used bitwise arithmetic for all genotype graph construction and manipulation routines which resulted in ˜100x speed-ups in some cases.
- We defined more compact data structures to store genotype graphs in memory which resulted in a >3x decrease in memory usage.
- We optimized the code for HMM computations to make it cache friendly and vectorised. This speeded up the HMM computations by ˜3x.

### Haplotype selection

A common approach to further speed up the sampling from P(D|H) is to only use a subset of K conditioning haplotypes instead of the full available set of 2N-2 haplotypes (with K << 2N-2). In SHAPEIT2^19^, we define the conditioning set as the K haplotypes minimizing the Hamming distance with the current estimate D = (h_1_, h_2_). In practice, this requires a quadratic scan of the haplotype data in O(LN^2^) that becomes prohibitive when facing large sample sizes (e.g. N > 10,000). In SHAPEIT3^5^, we improved this by maintaining a clustering of the haplotypes so that Hamming distance minimization is only performed on the haplotypes (typically 4,000) belonging to the same clusters than D. This approach exhibits close-to-linear running times in O(LNlogN). In SHAPEIT4, we propose a fully linear approach in O(LN) to assemble a conditioning set of K haplotypes that share long matches with D. To do so, we build a Positional Burrow Wheeler Transform (PBWT) of the full set of 2N haplotypes before each MCMC iteration (so 15 times total) by running the *algorithm 2* from *Durbin, 2014*^17^. This algorithm is very fast and only requires a single O(LN) sweep through the data. This gives prefix and divergence arrays that we store every 8 variants by default to reduce memory usage. A prefix array is a permutation vector of the original haplotype indexes in which haplotypes sharing long prefixes (i.e. matches) are sorted next to each other. A divergence array specifies the starts of the matches. We extended the algorithm to also give a third array relating original haplotype indexes to permuted ones so that any haplotype can be located in the prefix or divergence arrays in constant time. Given these PBWT arrays, we can then update the haplotypes for G as follows:

- We identify the P haplotypes sharing the longest prefixes with h_1_ and h_2_ at each stored variants. By definition, these haplotypes are sorted next to h_1_ and h_2_ in the prefix arrays and the procedure has therefore running times proportional to O(L).
- We collapse the 2LP/8 resulting haplotypes into a list of K distinct haplotypes (i.e. we remove duplicates). Long matches involve that the same haplotypes will be reported across multiple variants and therefore fewer distinct haplotypes once collapsed.
- We run the LSM onto the resulting conditioning set to get new estimates for G.

Using the same set of conditioning haplotypes across entire chromosomes is inefficient since some of them would only be informative at particular genomic regions. To account for this, we implemented the phasing procedure described above in a sliding window of size W (W=2Mb by default) similarly to what has been already done in SHAPEIT2. Overall, this gives a procedure that has multiple interesting properties. First, it provides informative haplotypes for the LSM as they are guaranteed to share long matches with D in the region of interest. Second, it is fast since building the PBWT and finding matches are done in running times proportional to O(LN). Finally, it gives a number of conditioning haplotypes that varies depending on the matches found in the PBWT: longer matches involve fewer distinct haplotypes and therefore smaller K. This implies that the size of the LSM (i) adapts to the local level of relatedness between individuals, (ii) shrinks as the MCMC converges and (iii) gets smaller for large sample sizes (i.e. sub-linearity with sample size).

### MCMC Iteration scheme

In SHAPEIT2 and 3, we start from a random haplotype resolution for all individuals and then perform 35 MCMC iterations to converge towards a reasonable haplotype resolution. In SHAPEIT4, we implemented three new features to improve the mixing of the MCMC:

- *Initialization*. We use a ‘quick-and-dirty’ initialization of the haplotypes so that the MCMC does not start from a random resolution. To do so, we use a simplified version of the phasing approach implemented in the PBWT software package^29^ which provides initial haplotype estimates very quickly in O(NL) with switch error rates in the range of 8% to 10% in our data. Briefly, we implemented a recursive procedure: at a given variant site l, we build a vector of all 2N alleles carried by the N individuals and set as missing all alleles at heterozygous/missing genotypes. Then, we infer a missing allele in this vector by copying from one of the two non-missing alleles carried by surrounding haplotypes in the PBWT indexes at variant l-1, making sure that the inferred allele is consistent with the original genotype. We iterate this update until no more missing data remains, then build the PBWT indexes for this variant l and move to the next one l+1.
- *IBD2 protection*. We designed an approach to prevent SHAPEIT4 to copy haplotypes across individuals sharing both of their haplotypes identical-by-descent (i.e. IBD2). This typically happens in the case of siblings and constitutes a ‘converge trap’ that can really hurt accuracy (the two individuals just copy their haplotypes without making any updates). To do so, we extended the algorithm 3 of Durbin, 2014^17^ to deal with the tri-allelic nature of genotype data and report genotype matches between individuals that are larger than W Mb (i.e. the sliding window). We then use the reported matches to define local constraints that we account for when building the conditioning sets of haplotypes. By constraint, we mean here triplets (i, j, w) where i and j are two individuals that are IBD2 in window w.
- *Specialized iterations*. We designed three different types of iterations to help MCMC mixing. A burn-in iteration (b) uses transition probabilities in the genotype graphs to sample new pairs of haplotypes. A pruning iteration (p) uses transition probabilities for sampling and also for trimming unlikely haplotype configurations in the genotype graphs by merging consecutive segments. Of note, this pruning iteration differs from previous SHAPEIT versions: instead of doing a single pruning stage made of multiple iterations (=8), we perform multiple stages of pruning (by default 3), each made of a single iteration. Finally, a main iteration (m) samples haplotypes and stores transition probabilities so that they can be averaged at the end of the run to produce final estimates. The user can specify any sequence of iterations and the default is 5b, 1p, 1b, 1p, 1b, 1p, 5m which we find to perform well.

As a consequence of these three features, SHAPEIT4 needs a small number of MCMC iterations to reach good level of convergence: by default, it only performs 15 iterations as opposed to previous versions that required 35 iterations (2.3x more iterations).

### Reference panel

SHAPEIT4 can borrow information from large reference panel of haplotypes which proves to be particularly useful when phasing only few individuals. To achieve this, SHAPEIT4 simply considers the additional reference haplotypes when building or updating the PBWT to allow conditioning haplotypes to also originate from the reference panel. Besides this, the iteration scheme remains remarkably identical to the algorithm described above. Of note, SHAPEIT4 only retains variants in the overlap between the main and the reference panel.

### Haplotype scaffold

SHAPEIT4 also allows for some heterozygous genotypes to be phased a priori. This approach has been previously introduced in SHAPEIT2 in the context of the 1000 Genomes project to perform genotype calling from low coverage sequencing data^19^. This ‘scaffold’ of pre-phased heterozygous genotypes can originate for instance from pedigrees in which many of the children’ genotypes can be accurately phased using Mendel inheritance logic. In practice, we implemented this functionality by simply pruning out all haplotype configurations in the genotype graphs that are inconsistent with the available scaffold of haplotypes (Supplementary Figure 5). Of note, SHAPEIT4 does not have any requirements in terms of variant overlap between the main genotype data and the scaffold haplotype data and therefore allows the latter to be derived from SNP array data while the former from high coverage sequencing data. In the context of this work, we demonstrate the potential of this approach by deriving scaffolds from either trios or massive reference panels.

### Phase informative reads

SHAPEIT4 allows haplotype estimation to be informed by sequencing reads overlapping multiple heterozygous genotypes. Haplotype assembly methods such as Whatshap^13^ or HapCut^14^ are very efficient at regrouping heterozygous genotypes within phase sets: groups of nearby genotypes for which the phase is inferred from sequencing reads that overlap them. In practice, SHAPEIT4 accommodates phase sets in a probabilistic manner (Supplementary Figure 5). When sampling new haplotypes for G, it assumes an error model in which paths in the genotype graphs that are consistent with the phase sets receive more weight than paths that are not. In other words, SHAPEIT4 samples haplotypes from the joint distribution P(D|H, R) ˜ P(D|H) x P(D|R) where R represents the available phase sets for G. The distribution P(D|R) is controlled by a parameter that defines the expected error rate in the phase sets. By default, we assume an error rate of 0.0001 and used this value in all reported results. As a consequence, the sampled haplotypes D for G are not necessarily consistent with all the phase sets which allows correcting phasing errors in sequencing reads when phase sets are too discordant with population based phasing.

### I/O interface

All the input and output interface implemented in SHAPEIT4 is built on the High-Throughput Sequencing library (HTSlib^30,31^) so that genotype and haplotype data is read and written in either Variant Call Format (VCF) or its binary version; the BCF format. This has multiple benefits. All data management on input and output files can be done using standard tools such as bcftools^31^. Using the BCF format significantly speeds up I/O operations. VCF/BCF formats natively define phase sets which facilitates the integration of phase information contained in sequencing reads in SHAPEIT4. HTSlib allows reading simultaneously multiple VCF/BCF files which facilitates complex SHAPEIT4 runs combining for instance genotype data with phase sets, a reference panel of haplotypes and a scaffold of haplotypes.

### The UKB datasets

We generated genotype data sets with large sample sizes from the full release of the UK biobank (UKB) data set containing 488,377 individuals genotyped by SNP arrays at 805,426 genetic variants. To do so, we proceeded according to the five following steps. First, we filtered out all genetic variants with more than 5% missing genotypes. Second, we extracted data only for chromosome 20, resulting in 18477 variants in total. Third, we build trio candidates (two parents and 1 child) from the pairwise kinship and IBS0 estimates between individuals that have been measured as part of the UKB study^2^ and took the first 500 trios minimizing Mendel inconsistencies between parents and children genotypes. The 500 children constitute high quality validation data since their haplotypes can be almost entirely resolved using Mendel inheritance logic. Fourth, we merged these 500 individuals with multiple random subsets of other UKB individuals in order to build 11 genotype data sets comprising between 500 and 400,000 individuals in total. Finally, we remove all individuals containing more than 5% missing genotypes in each data set. This gave us 11 data sets comprising 499, 995, 1,997, 4,973, 9,939, 19,894, 49,752, 99,452, 198,894, 298,383 and 397,839 individuals for which genotype data is available at 18,477 variants. To mimic phasing from large reference panels, we used the best haplotype estimates obtained for each sample size from which we removed the haplotypes of the 500 children. This resulted in 10 reference panels comprising 992, 2996, 8948, 18,880, 38,790, 98,506, 197,906, 396,790, 595,768 and 794,680 haplotypes defined at 18477 variants. All reference panels have been compressed to speed up I/O of the downstream phasing runs: we used BCF formats for both SHAPEIT4 and Eagle2 (using bcftools v1.4) and bref3 format for Beagle5 (using bref3.28Sep18.793.jar from https://faculty.washington.edu/browning/beagle/). For the main panels that were phased against these reference panels, we used the 500 trio children that we merged with 4,500 and 49,500 other unrelated UKB samples. This resulted in 3 main panels with 500, 5,000 and 50,000 individuals with genotype data at 18,477 variants. Of note, all data management was done using bcftools v1.4^31^.

### Phasing runs on the UKB datasets

We phased the UKB datasets using SHAPEIT4 and three other widely used phasing methods: SHAPET3 (https://jmarchini.org/shapeit3/), Eagle v2.4 (https://github.com/poruloh/Eagle) and Beagle5 (https://faculty.washington.edu/browning/beagle/beagle.html). All runs were done on a RedHat server with Intel(R) Xeon(R) CPU E7-8870 v4 @ 2.10GHz (80 physical cores and 160 logical cores) and 3 Tb of RAM. Eagle v2.4 and Beagle5 have been run with default parameters. SHAPEIT3 has been run using the options --states 200, --cluster-size 4000 and --early-stopping. All methods were run using 10 threads to speed up computations. The reported running times and memory usages were obtained using the GNU time v1.9 command. The running times are computed as the sum of the User and System times and the memory usage as the Maximum resident set size. The error rates of each method was measured using the switch error rate (SER) on the 500 trio children. Specifically, we enumerated all heterozygous genotypes (i.e. hets) in the 500 trio children that can be phased using Mendel inheritance logic (i.e. no triple hets and no Mendel inconsistencies). Then, we computed the SER as the fraction of successive pairs of hets that are correctly phased over all possible pairs. Confidence intervals for the SER are defined as binomial 95% confidence intervals and were computed using the R/binconf of the R/Hmisc package^32^.

### The GIAB datasets

To assess performance of SHAPEIT4 on sequencing data, we use the high quality phased genotype data generated for the NA12878 individual by the Genome In A Bottle consortium^22^. In order to get population scale data, we merged the high coverage genotype data of the GIAB with 502 unrelated European individuals sequenced as part of the phase 3 of the 1000 Genomes project phase 3 (KGP3)^24^. To do so, we proceeded as follows. First, we only used variants on chromosome 20. Second, since NA12878 has also been sequenced in KGP3, we only retained variants for which GIAB and KGP3 genotypes are concordant. Of note, NA12878 is not included in the 502 KGP samples we used for merging. Third, we only kept variants that were phased by GIAB thanks to some family data (i.e. multiple sequenced individuals from the NA12878 family). Fourth, for all variants present in KGP3 and not in GIAB, we assume NA12878 to be homozygous reference allele. In total, this procedure gave us a data set on chromosome 20 comprising genotype data for 503 European individuals across 507181 genetic variants amongst which 478581 are SNPs (the rest being a mixture of short indels and large SVs). Of note, all data management was done using bcftools v1.4^31^. The GIAB consortium has generated sequence data using multiple sequencing technologies for NA12878. In the context of this work, we notably used: (a) SOLiD5500W, (b) Complete Genomics, (c) Illumina HiSeq, (d) PacBio and (e) 10x Genomics. Summary statistics for all this sequence data is given table 1. We used the 5 sets of sequencing data to phase as much as possible of the NA12878 genotype data. In practice, we used WhatsHap v0.15^13^ with default parameters to phase NA12878 using sequence data (a) to (d). The proportion of heterozygous genotypes being phased by each type of sequence data is shown in table 1. For 10x Genomics sequence data, we proceeded quite differently since this technology relies on barcoded reads and not on long reads. In this case, we used the set of heterozygous genotypes that can be phased using the Long Ranger method provided by 10x genomics. We have not run the method ourselves but instead used the Long Ranger outcome provided by the GIAB consortium. The outcome of WhatsHap or Long Ranger has been included in the VCF file as phase sets (i.e. field PS) so that SHAPEIT4 can leverage it.

### Scaffolding the GIAB data

We generated two haplotype scaffolds for the dataset described above either using family or large reference haplotype panel. For the first scaffold, we used genotype data derived from Illumina OMNI2.5M for a larger set of individuals containing multiple trios and duos. Out of the 503 individuals with sequence data, 190 of them (˜37.8%) are present in the OMNI data together with parents or children so that we could fix the phase at 6.59% of the heterozygous genotypes in the sequence data using Mendel inheritance logic. For the second scaffold, we phased the 16805 variants in the overlap with UKB data on chromosome 20 using 795678 UKB haplotypes as reference panel. This allowed fixing the phase at 6.25% of the heterozygous genotypes in the sequence data.

### Phasing runs on the GIAB datasets

Using SHAPEIT4, we phased the 503 individuals with sequence data across all possible combinations of 6 types of sequencing reads (i.e. no reads, SOLiD5500W, Complete Genomics, Illumina HiSeq, PacBio and 10x Genomics) and 3 scaffolds (no scaffold, OMNI and UKB scaffolds). Each run was repeated 5 times using different random number generator seeds to assess variability of the results. Accuracy of phasing was measured using the switch error rate as described above on NA12878 using the phase provided by GIAB as reference.

### Code availability

SHAPEIT4 is available on the GitHub webpage (https://odelaneau.github.io/shapeit4/). The code is licenced under the MIT licence.

### Data availability

We used the full release of the UK biobank (http://www.ukbiobank.ac.uk/). This work was conducted under UK Biobank project 35520. We used the following release of the GIAB (ftp://ftp-trace.ncbi.nlm.nih.gov/giab/) and this release of the phase 3 of the 1000 Genomes project (ftp://ftp.1000genomes.ebi.ac.uk). We downloaded this version of the larger set of 1000 Genomes samples genotyped on Illumina OMNI2.5M (ftp://ftp.1000genomes.ebi.ac.uk). The sequencing data we used for GIAB are available from these locations: PacBio (<u>BAM</u>), Illumina HiSeq (<u>BAM</u>), SOLiD5500W (<u>BAM</u>), Complete Genomics (<u>BAM</u>) and 10X Genomics VCF populated with with PS field (<u>BAM</u>).

**Supplementary Figure 1:**
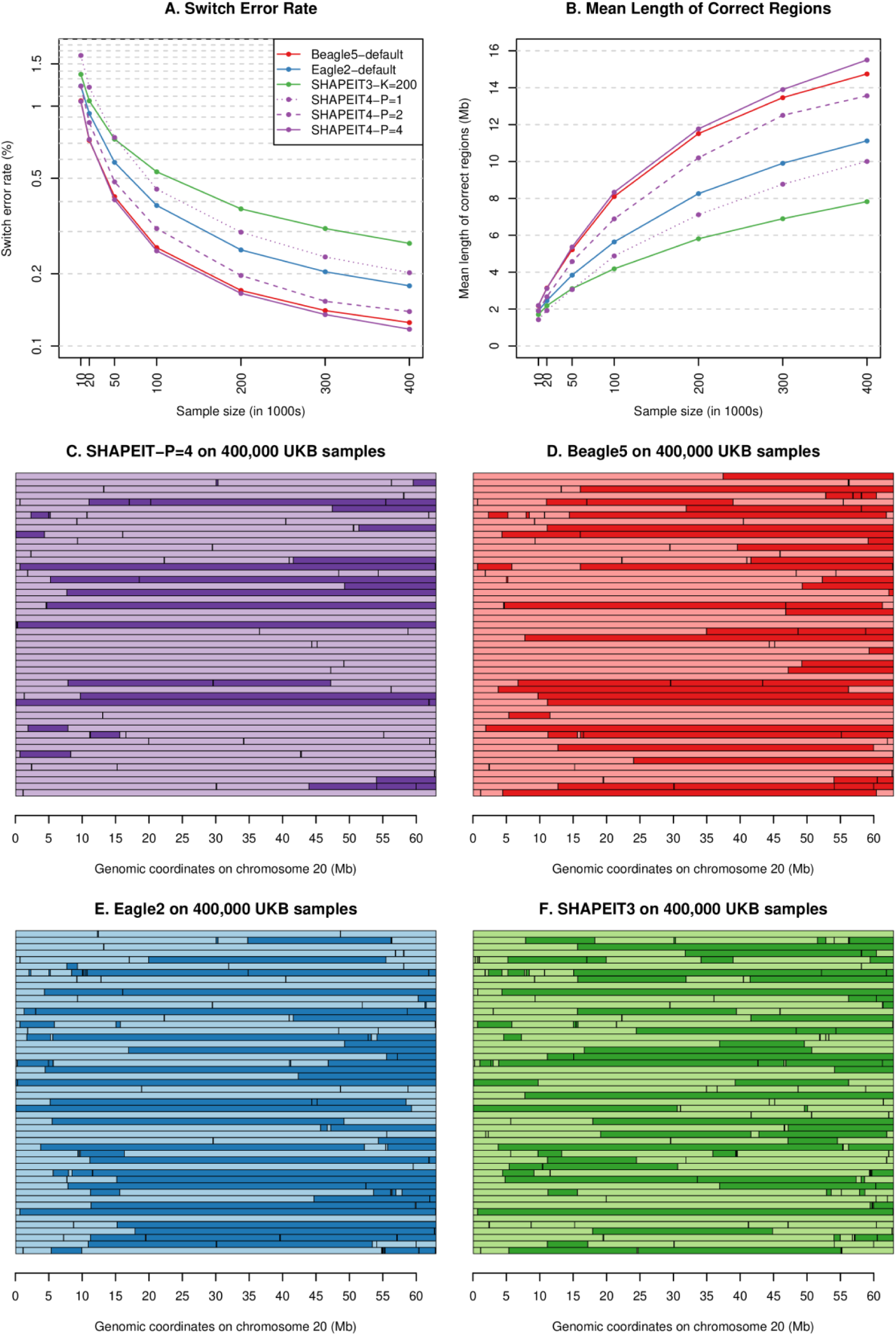
Switch errors in UKB. Switch error rates (**A**) and average genomic distances between switch errors (i.e. size of the regions being correctly phased; **B**) as a function of sample size and tested method. Genomic locations of the switch errors on chromosome 20 across 50 UKB samples used for validation when inferred using SHAPEIT4-P=4 (**C**), Beagle5 (**D**), Eagle2 (**E**) an SHAPEIT3 (**F**) on 400,00UKB samples. Any switch between light and dark colour stands for a switch error.

**Supplementary Figure 2:**
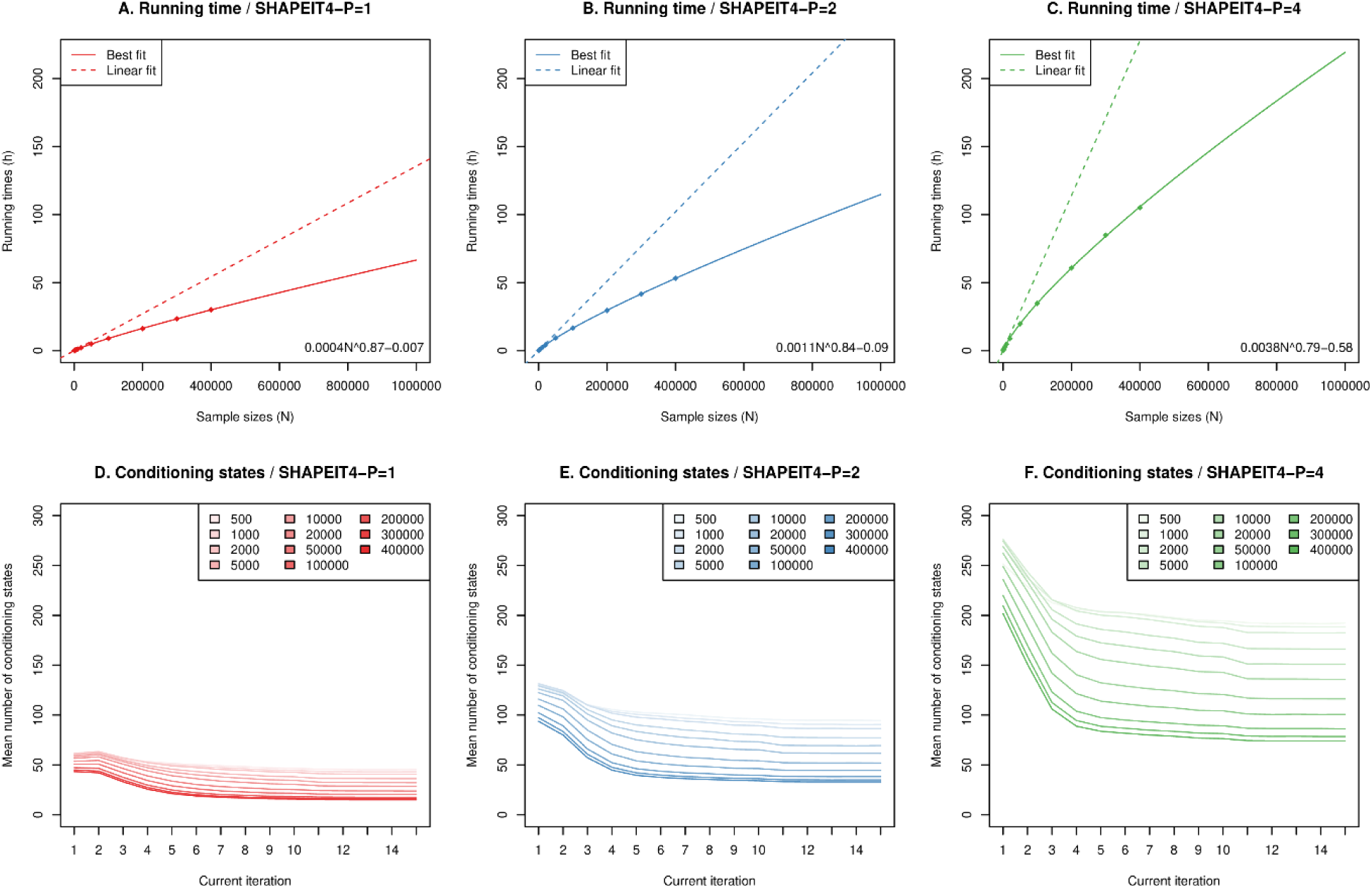
SHAPEIT4 complexity with sample size. Running times as a function of UKB sample size for SHAPEIT4 when run with P=1 (**A**), P=2 (**B**) and P=4 (**C**). The dots show the measured running times. The plain lines show the fitted function (detailed on the bottom right corner) on the available data points. The dashed lines show the linear fit obtained when fitted to the first two data points. Variation of the mean numbers of conditioning haplotypes used by SHAPEIT4 as a function of the iteration (from 0 to 15) for each of the UKB data sets tested (from 500 to 400,000 individuals) and across multiple SHAPEIT4 parameter values (**D-F**).

**Supplementary Figure 3:**
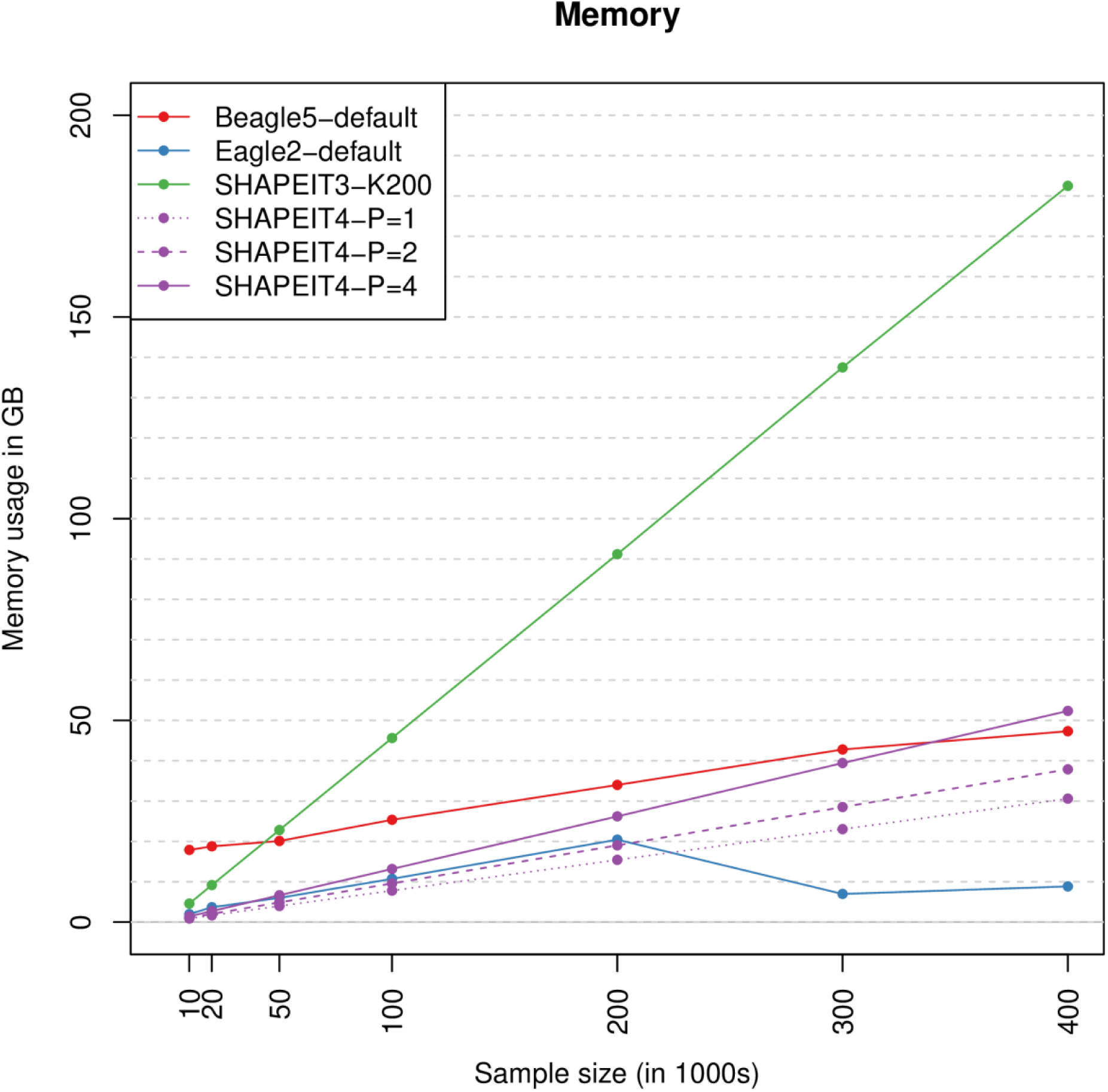
Memory usage. Memory usage in GigaBytes as measured by the Linux time command as a function of the sample size and the method tested.

**Supplementary Figure 4:**
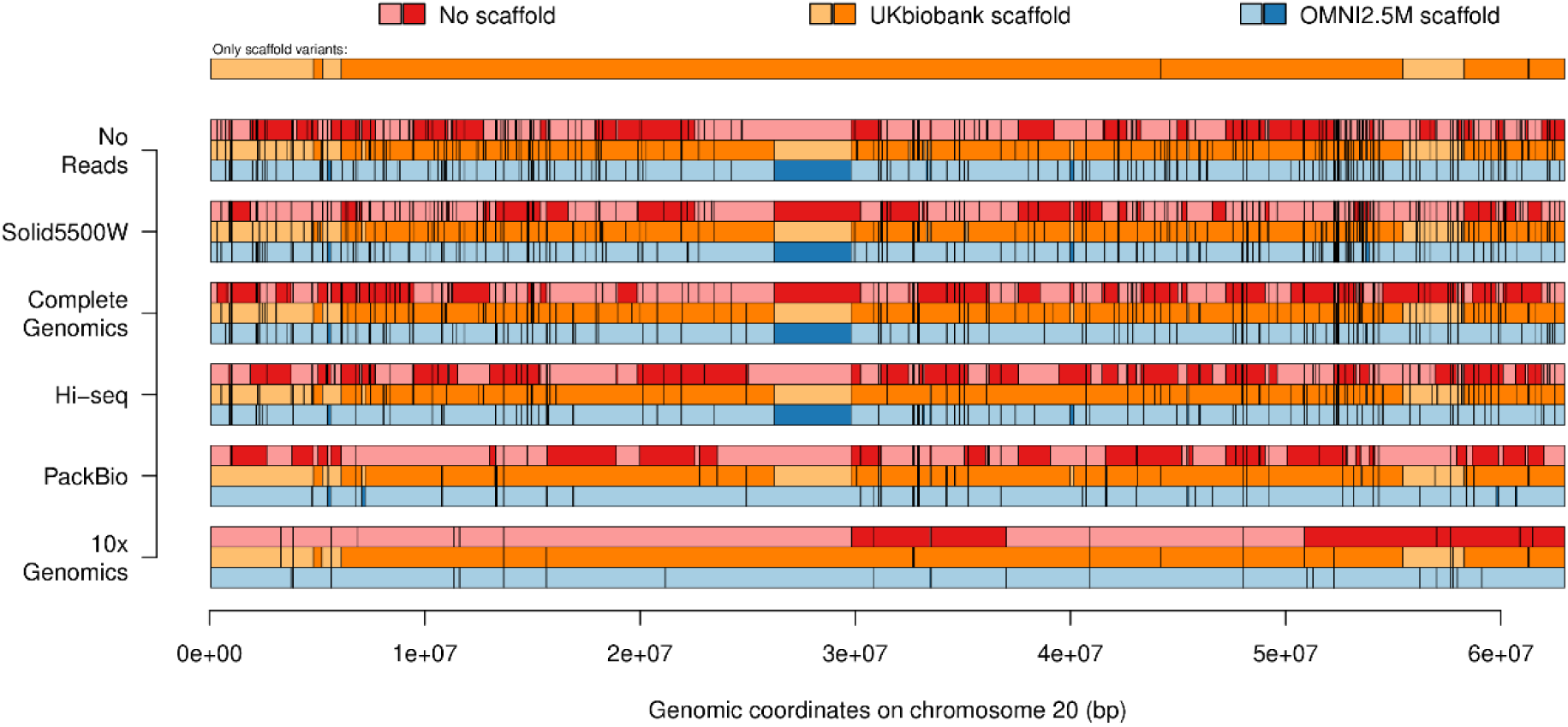
Switch errors in GIAB. Genomic locations of the switch errors on chromosome 20 for each one of the 18 tested configurations. Switches between dark and light colour represent switch errors. The top row shows the switch errors only between variants in the overlap with UKB when NA12878 is phased against UKB (i.e. accuracy of the UKB scaffold).

**Supplementary Figure 5:**
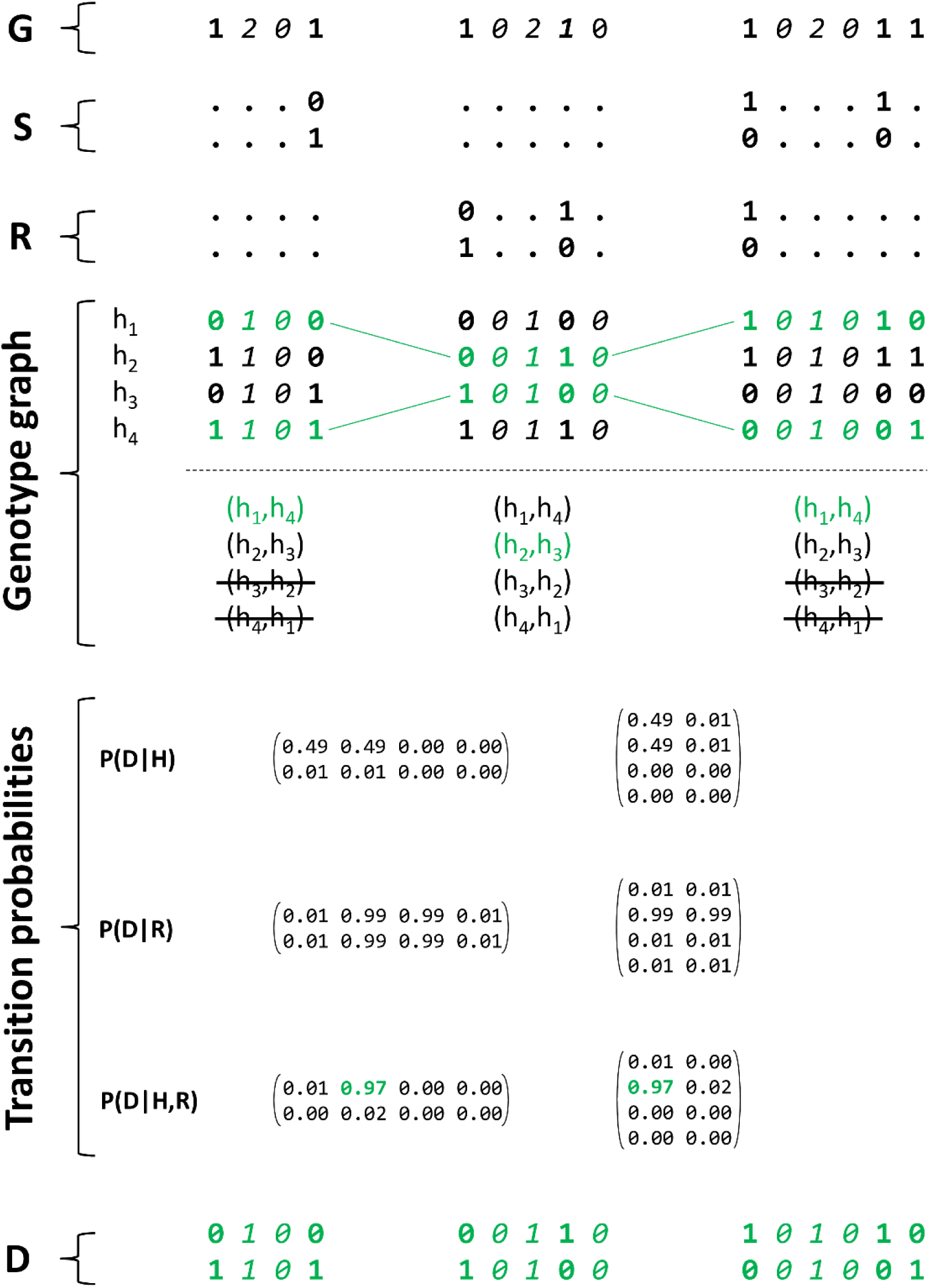
Overview of the SHAPEIT4 genotype graph structure. From top to bottom. (1) The genotype data **G** available for an individual with 0, 1 and 2 corresponding to homozygous reference, heterozygous and homozygous alternative genotypes. Non heterozygous genotypes are italicized. (2) The haplotype scaffold **S** available for this individual with three pre-phased heterozygous genotypes. (3) The phase set **R** available for this individual at three heterozygous genotypes. (4) The genotype graph structure for **G** when assuming only two heterozygous genotypes per segment (it is three in SHAPEIT4). On the top panel, the four possible haplotypes per segment. On the bottom panel, the pairs of haplotypes consistent with both **G** and **S**. (5) Examples of transition probabilities between the three segments when derived using the Li and Stephens model, P(D|H) and when derived from the phase sets P(D|R). Assuming an error rate of 0.01, transitions not consistent with the phase sets are assigned a probability of 0.01 and those that are a probability of 0.99. The last sets of transition probabilities P(D|H,R) are those from which SHAPEIT4 samples haplotypes for G. An example of sampled haplotypes D for G is shown in green.

